# Nature-based solutions in Australia: a systematic quantitative literature review of terms, application and policy relevance

**DOI:** 10.1101/2023.05.11.538642

**Authors:** Dan Zhu, Lily Fraser, Dave Kendal, Yue Zhang, Emily J Flies

## Abstract

Nature-based Solutions (NbS) are emerging as an approach to sustainable environmental management and addressing environmental and social issues in ways that benefit human well-being and biodiversity. NbS have been applied to social-environmental challenges such as climate change and urbanization, but with diverse conceptualisations and applications that may impact their effectiveness and broader uptake. Much of the literature and implementation of NbS has emerged from Europe and though NbS use is rising in Australia, the context is unclear. This systematic quantitative literature review aims to understand Nature-based Solutions in an Australian context.

Here we explore the meaning and practical uses of NbS in Australia, through three research questions: In Australia, 1) what is meant by the term ‘nature-based solutions’? 2) what socio-ecological challenges do NbS aim to address and how? 3) are there gaps in NbS research and policy application that are hindering uptake of NbS approaches?

We show that in Australia, local governments are using NbS in urban planning to address the compounding challenges brought on by climate change in the human-environment interfaces. However, there is no consensus on NbS definitions and approaches, research is focussed on urban areas and problems, and NbS implementation follows a bottom-up, localised pattern without an integrated policy framework. Based on these findings, we provide recommendations for improving the implementation of NbS in Australia including: 1) a consistency of NbS definition and awareness of NbS approaches; 2) interdisciplinary and interdepartmental collaboration on NbS methods and effectiveness and; 3) an integrated policy framework supporting NbS nationwide.

## 1 Introduction

Climate change, biodiversity loss and environmental degradation bring enormous challenges to nature and human society; sea level rise, droughts, bushfires, soil erosion, decreasing biodiversity, and flooding cause trillions of dollars in loss of crops, forests and urban infrastructure (McMichael et al. 2008; Wheeler and von Braun 2013; Bellard et al. 2012). The interconnected causes and impacts of environmental change further increase food insecurities and pandemic outbreaks, which pose risks to global societies (Tirado et al. 2010; Daszak et al. 2001). Mitigating and adapting to these impacts effectively has become a primary challenge for global decision-makers and societies.

Nature-based Solutions (NbS) are increasingly seen as an effective tool for dealing with socio-environmental challenges, building resilience and driving sustainability transitions (Fastenrath, Bush & Coenen 2020). A commonly used definition for NbS has been the International Union for Conservation of Nature (IUCN) definition: “actions to protect, sustainably manage and restore natural and modified ecosystems in ways that address societal challenges effectively and adaptively, to provide both human well-being and biodiversity benefits” (IUCN 2022). This definition builds on the recognition that society relies on healthy and functioning ecosystems to thrive. The United Nations Environment Assembly (UNEA) recently adopted a multilaterally agreed-upon definition similar to the IUNC definition: ‘actions to protect, conserve, restore, sustainably use and manage natural or modified terrestrial, freshwater, coastal and marine ecosystems, which address social, economic and environmental challenges effectively and adaptively, while simultaneously providing human well-being, ecosystem services and resilience and biodiversity benefits.’ (UNEA 2022)

NbS in cities and in rural areas can be used to address major societal challenges like climate change, food and water security, disaster risk reduction, and health while supporting biodiversity, wellbeing and sustainable development (IUCN 2022). In the past five years, urban forests, green roofs and walls, parks, river restoration and many other forms of NbS have come into the mainstream of urban planning and policy-making (Fastenrath, Bush & Coenen 2020). The benefits of NbS for people and environments have made them of increasing interest for government strategies and policies (Bayulken, Huisingh &Fisher 2021).

Australia is on the frontline of climate change impacts, experiencing the predicted changes to climate and increase in severe weather with considerable damage to nature and society. For example, the 2019-2020 bushfires burned more than 46 million acres of land and caused the loss of 3,500 homes, 34 human lives (CDP 2020) and hundreds of millions of animal lives (The University of Sydney 2020). Many Australian cities are coastal and are threatened by rising sea levels, heatwaves are causing deaths and reducing the liveability of already hot cities, and increasingly frequent and severe flooding affects thousands of Australian homes (AAS, 2021). Australia is also facing a biodiversity crisis with extremely high rates of species extinction and continuing loss of habitat, entangled with the impacts of climate change. NbS are an important tool in addressing climate change and have important benefits for biodiversity. Australia’s *Strategy for Nature 2019-2030* describes NbS as “critical to build the resilience of our unique nature” (Commonwealth of Australia 2019).

Examining the practice of and barriers to the use of NbS in Australia is essential for a better implementation and effective sustainable development. To facilitate uptake of NbS in policy and practice in Australia, this review examines four research questions:

1. What is meant by the term ‘nature-based solutions’ in Australia?
2. In Australia, what problems does research on NbS aim to address and how?
3. What are the research gaps and barriers to greater integration of NbS into policy and practice in Australia?

These questions are explored through a systematic review of the literature. Figure. 1 shows the conceptual model of this study around which the paper is structured. By analysing existing literature systematically, we are able to produce a picture of what NbS means in Australia, where the research and policy gaps are and make recommendations of how to support greater uptake of NbS in Australia.

**Fig.1.**
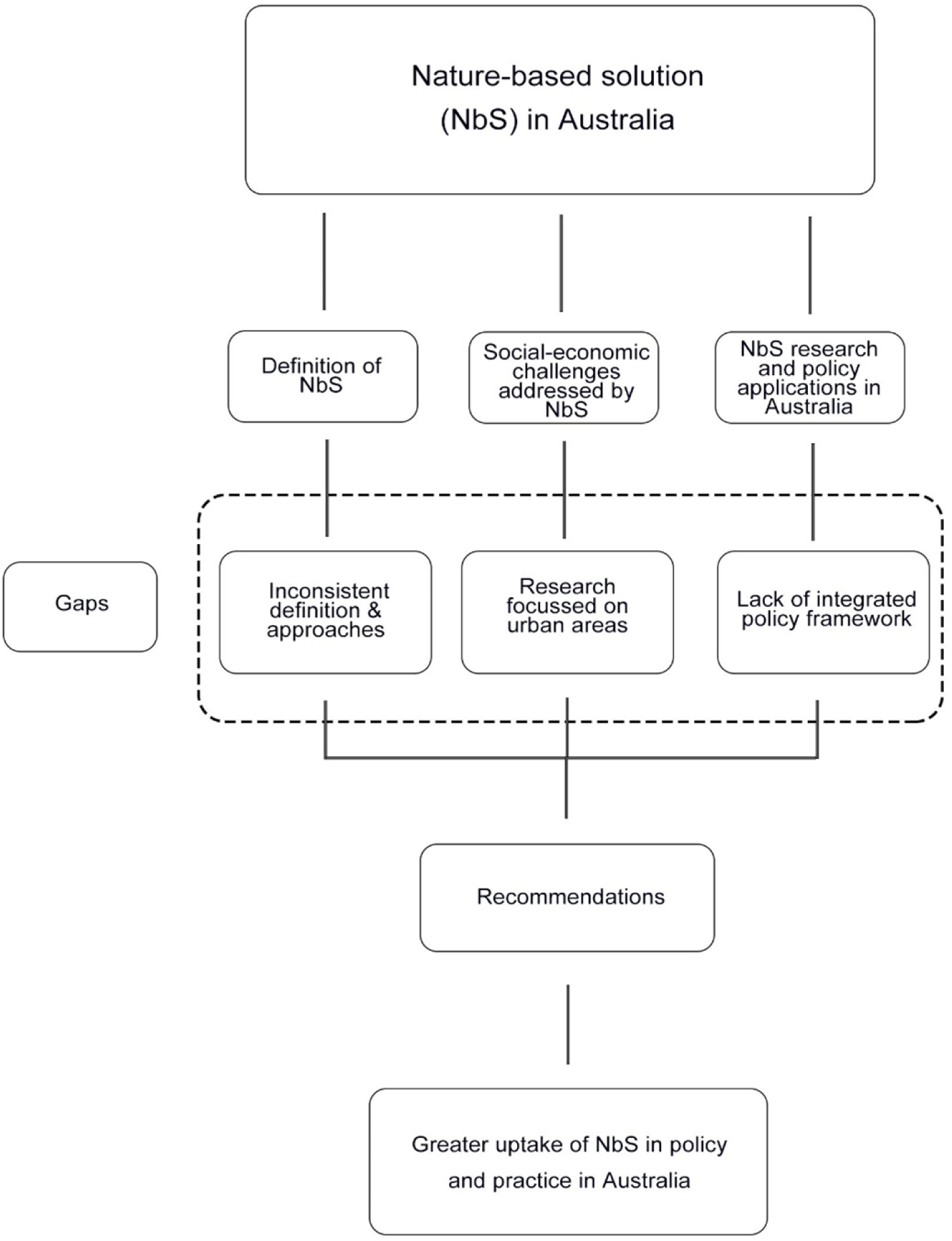
Conceptual model of understanding NbS in Australia around which this literature review is structured

## 2 Methods

The systematic quantitative literature review method applied in this study allows researchers to examine existing studies to identify research trends and gaps in a systematic and replicable way (Pickering & Byrne 2014). We used the Preferred Reporting Items for Systematic reviews and Meta-Analyses (PRISMA) approach (Figure 2) as a transparent, precise, and complete record of the paper selection process for this review (Page et al. 2021).

**Fig. 2.**
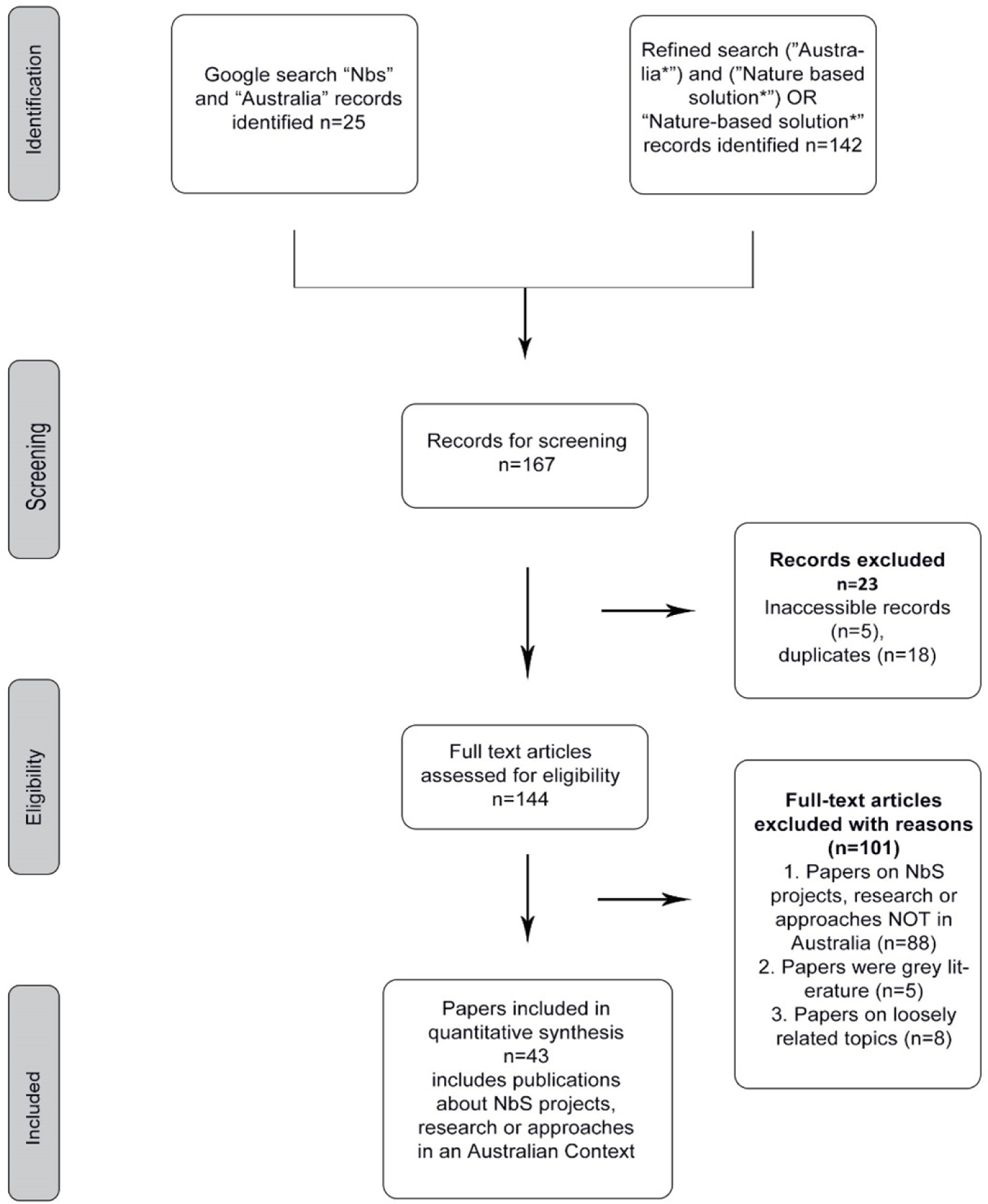
A PRISMA diagram of the method and process used in this literature review

### 2.1 Search strategy

The systematic search of Web of Science was conducted on 6 May 2022 and used the search terms: ***(“Australia*”) AND (“Nature based solutions*” OR “Nature-based solutions*”)*** in “all fields” (including publication titles, abstracts, keywords and other fields such as publication name and author affiliation), retrieving both articles about NbS in Australia, and articles about NbS by authors at Australian institutions. These results were combined with additional relevant articles from a search of Google Scholar. To contextualize Australian NbS research against the global literature published on the Web of Science, “Australia’’ was removed from the search terms ***(“Nature based solution*” OR “Nature-based Solution*”)*.**

A set of inclusion and exclusion criteria were used to screen articles (Table 1). During the screening and eligibility stages, Rayyan.ai, an online AI software, was used as a collaboration tool (Ouzzani et al. 2016). This facilitated real-time changes by group members and online discussions about eligibility to improve the final paper selection.

**Table 1.** Inclusion and exclusion criteria are based on methods recommended by Pickering & Byrne 2014 and the University of Melbourne (2022)

| **Inclusion** | **Exclusion** |
| --- | --- |
| All areas, fields or disciplines using NbS in Australia | NbS projects, research or approaches NOT based in Australia |
| Articles discussing projects, research or approaches to NbS in Australia | Grey Literature, book chapters, presentation abstracts and conference papers |
| Peer reviewed articles | Loosely related information |
|  | Duplicate articles |

### 2.2 Data extraction and analysis

The final articles included for quantitative analysis (Appendix 1) were recorded in the reference management system Zotero. An Excel spreadsheet, referred to as the “database”, was created containing all articles. Metadata from each article was also coded into the database, including study location, year of publication, publication title. Clarivate InCites was used to analyse the subject area of the journals that the included articles were published in. Descriptive codes were used to categories articles based on the methods used and policy referred to. Articles were categorised as using qualitative, quantitative or mixed methods. Qualitative methods included interviews, opinion papers and reviews. Quantitative methods included questionnaires, and models while mixed method studies included both qualitative and quantitative methods. Articles were also coded to indicate whether they used case studies, document analysis and/or modelling approaches. Any reference to NbS policy was coded by level of government - local, state, federal or international.

Coding criteria were developed using definitions, key attributes and conceptual ideas identified by group members during the scoping and screening phases. Articles were assigned to different members of the group who coded each using the full text. The main coding categories were derived from the research questions: 1) approaches to NbS, 2) problems being addressed by NbS 3) scale of implementation and 4) barriers and opportunities for NbS implementation. Each category was further subdivided into specific sub-categories that reflected the terms that operationalised the category concepts. For example, in the problem category of ‘natural disasters’, an could use terms such as ‘bushfires’ or ‘floods’. Approaches were coded following the IUCN (2020) categories of (1) restoration or (2) protection, (3) issue-specific approaches (determined by site-specific natural and cultural contexts), (4) infrastructure related approaches (incorporating natural features into infrastructure planning and development), and (5) ecosystem management approaches (which see humans as key components of the ecosystem by integrating ecological, social and economic goals) (IUCN 2020, Table 2). Any other approaches which did not fit these categories were classified as “other terms”. As many articles discussed multiple approaches to NbS, the total number of papers that mention an approach was greater than the number of papers reviewed. The problems or issues that the NbS aimed to address were similarly coded into categories and sub-categories, scale was coded as local, regional (e.g. statewide) and national, and where barriers to implementing NbS policy were mentioned, these were also recorded.

**Table 2.**
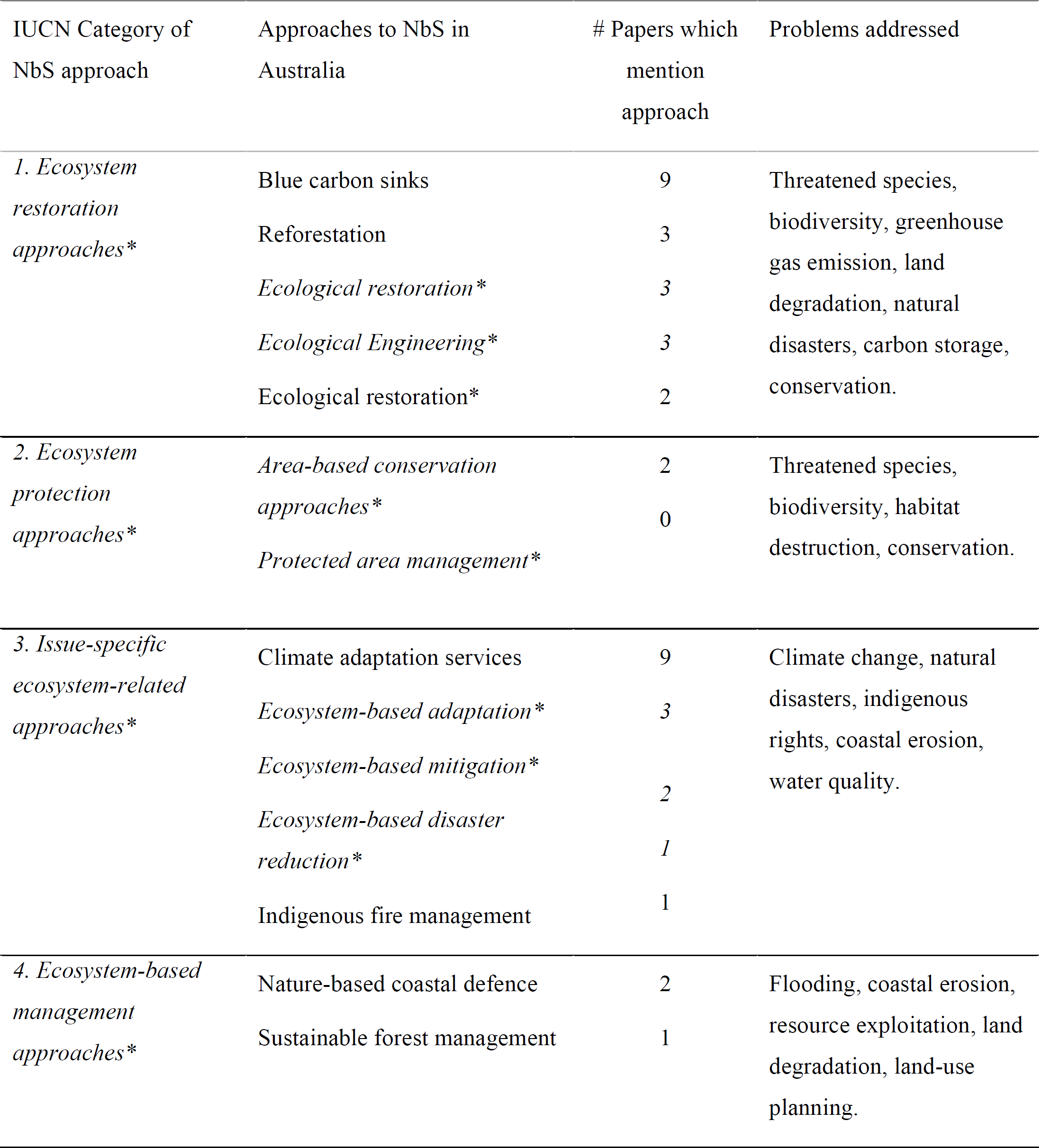
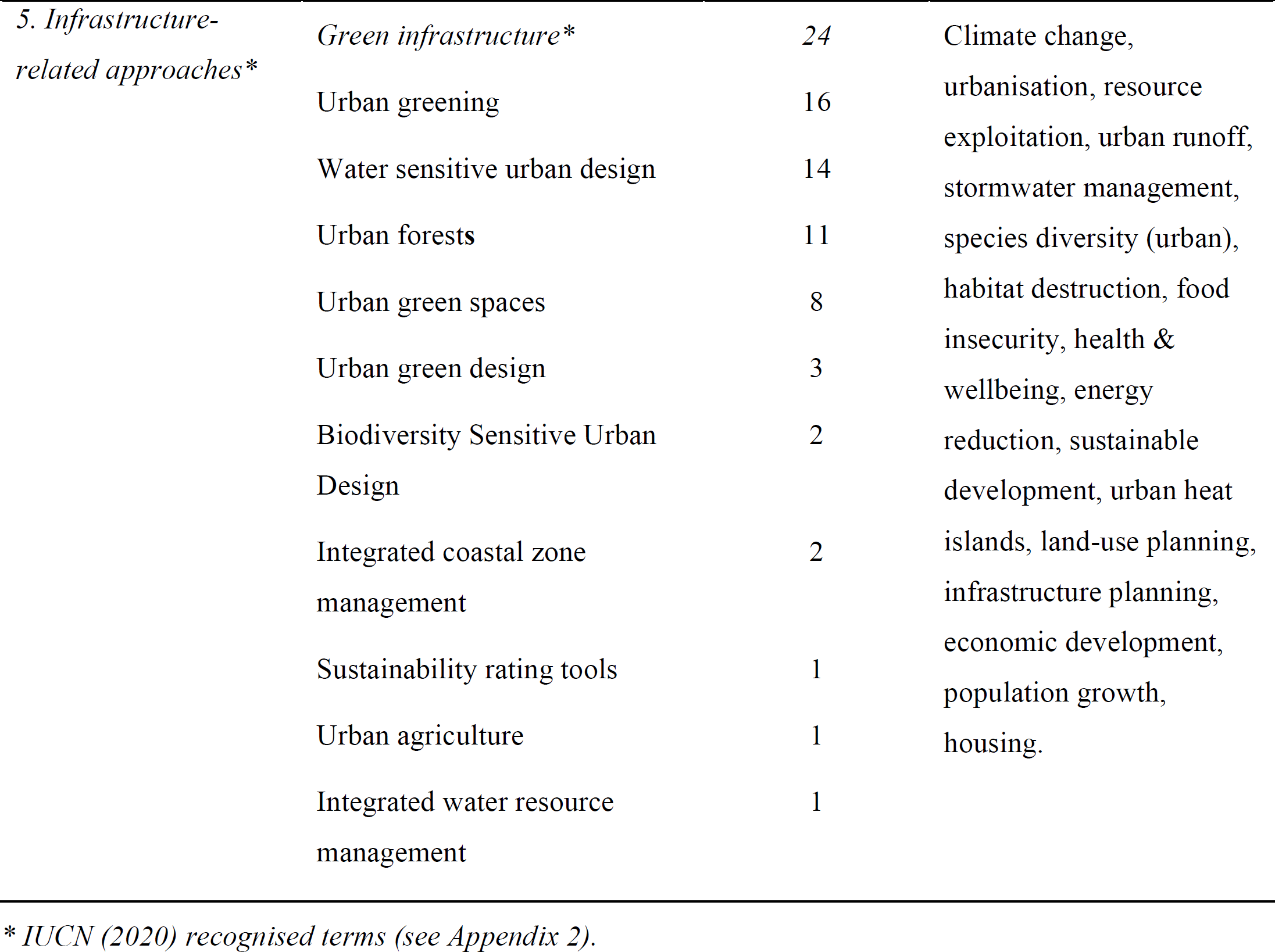
Summary of approaches and problems addressed by NbS in Australia.

## 3. Results

Our Web of Science search returned 142 documents with an additional 25 documents sourced from Google Scholar (n = 25). Screening by Rayyan.ai and group members identified duplicates (n=18) and inaccessible articles (n=5). The remaining articles (n=144) were then assessed against the inclusion/exclusion criteria (table 1). Articles were excluded where projects, research or approaches outside of Australia (n=88), grey literature (n=5) and on loosely related topics (n=8). A final list of 43 were included in the analysis.

### 3.1 Contextualising NbS in Australia

The global search (excluding the Australia term) returned 1,581 documents, indicating that less than 1% of the literature published on NbS in Web of Science is done in Australia or by Australian authors.

In the Australian context, there has been a steady increase of literature published on NbS in Australia (Figure 3). The first articles were published in 2017 (Greenway 2017). Over the 5 years from 2017 to 2021, there was a 2000% increase in the number of articles published annually.

**Fig. 3.**
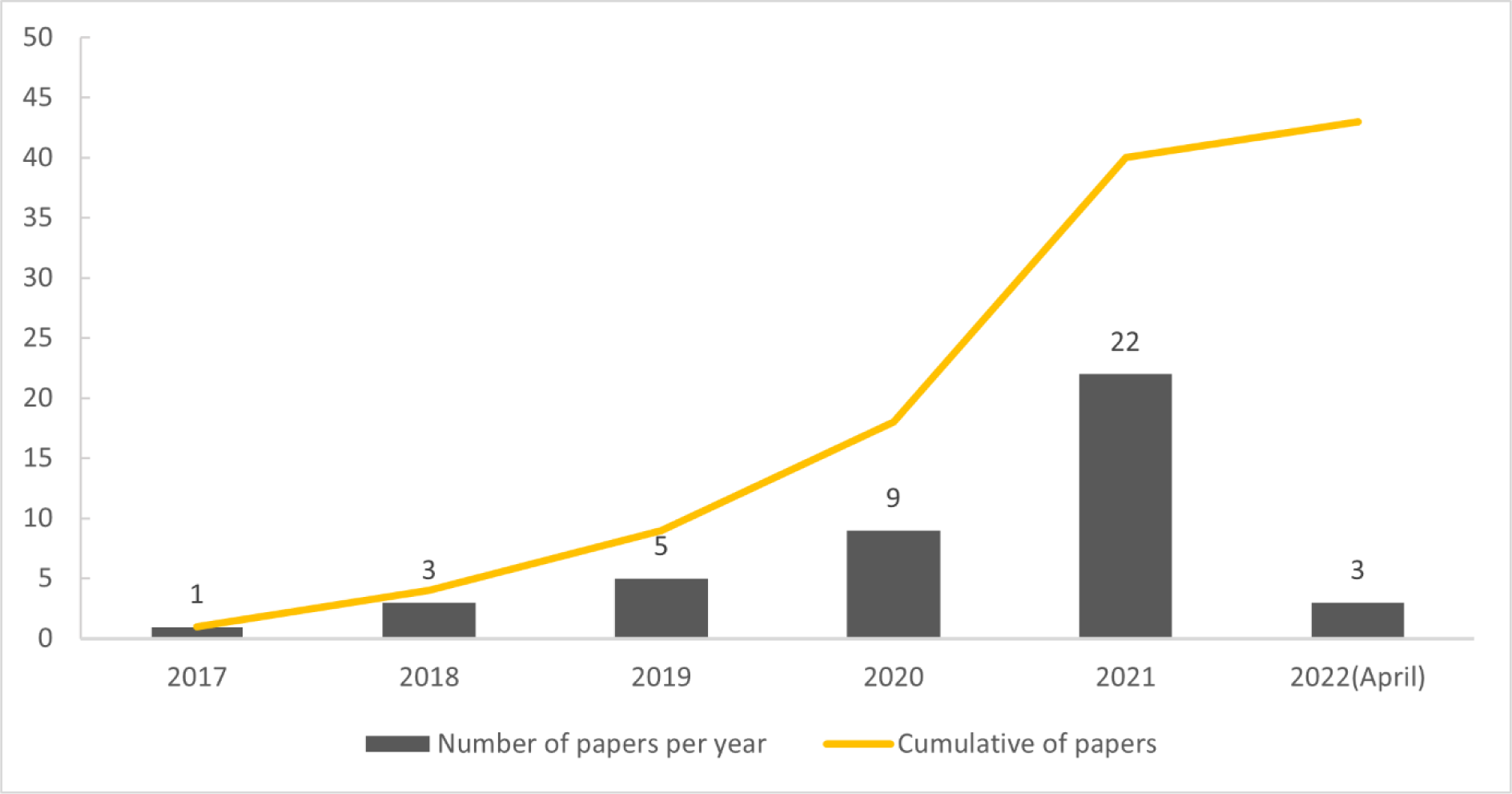
Number of papers published on NbS in the Australian context over time

Science and environment journals published the most articles on NbS (n=39). However, many articles were published in multi-disciplinary journals, with urban studies (n=21), planning (n=19), urban and policy (n=16) also being common.

### 3.2 Characteristics

The current Australian literature on NbS is largely based in major cities (n=39), and predominantly in Victoria (Figure 4). Specifically, NbS projects, strategies and approaches in Melbourne were mentioned the most frequently (n=25). The papers published by Morris et al. (2019), Ossola & Lin (2021), Rogers et al. (2020) were the only to refer to NbS outside of major cities, and Shumway et al. (2021) referred to NbS nationally.

**Fig. 4.**
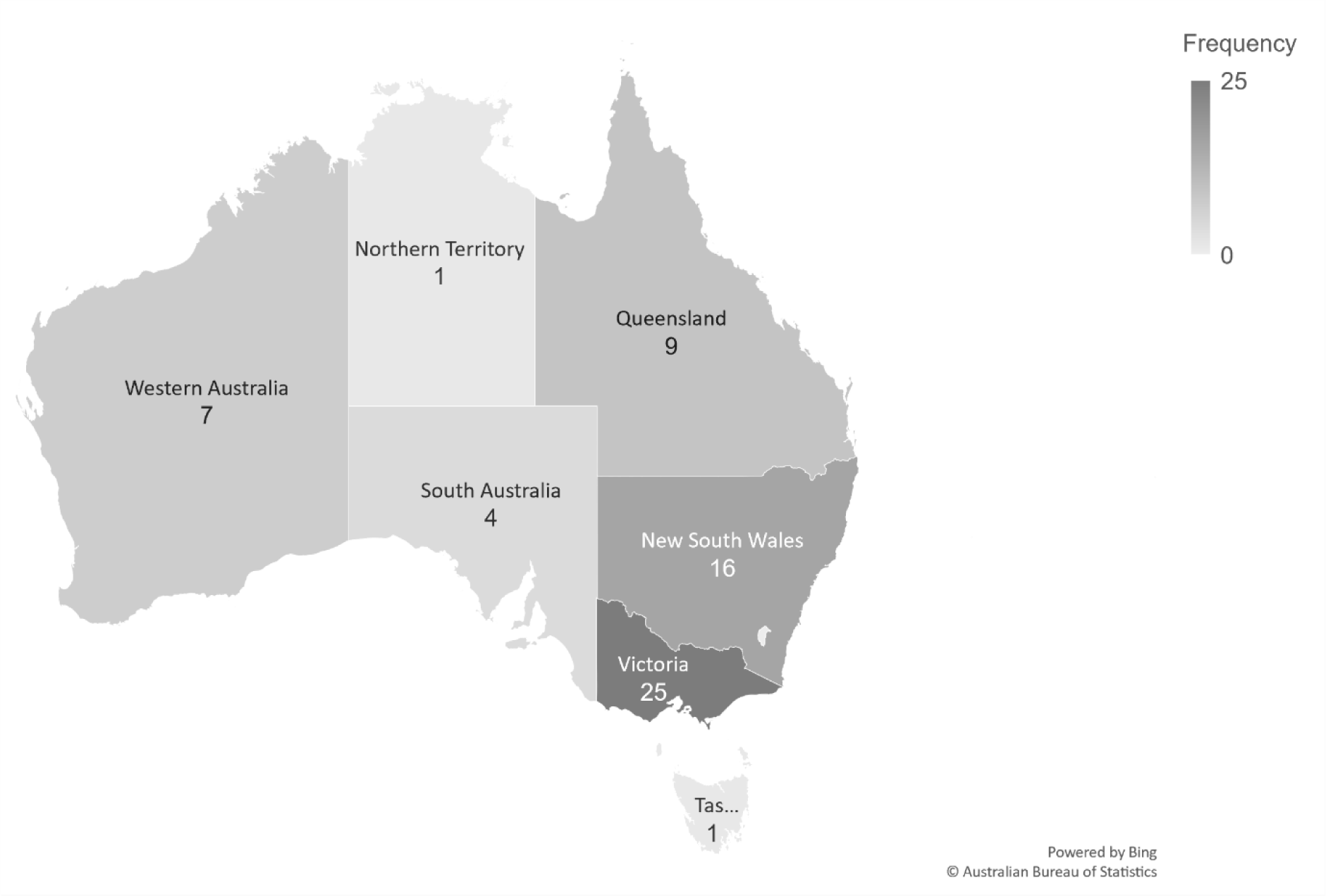
Map showing where NbS research, projects and approaches are occurring in Australia. State totals sum greater than the number of papers reviewed, as some papers mentioned more than one location.

There was a relatively even spread across study methodologies including: qualitative (n=15), quantitative (n=12) and mixed methods (n=16) studies. The most favoured research methods used in the literature were case studies (n=21), document analysis (n=13) and modelling (n=11). Note that these methods values sum to greater than the number of papers reviewed because many papers used multiple methods of data collection.

### 3.3 Definitions

In total, 63% of papers reviewed did not provide a definition of NbS (n=27). Of the papers which defined NbS (n=16), six used the IUCN definition and Ignatieva et al. (2020) used a definition provided by the European Commission. Other definitions (n=9) were often referenced from other papers published on NbS in the literature. The Oxford definition of NbS was searched for in all 43 papers, but none were found to use it (n=0).

### 3.3 Approaches to NbS

The IUCN approaches to NbS were mentioned a total of 52 times. However, non-IUCN terms (i.e. ‘other terms’) were mentioned more frequently than IUCN approaches (n=71) demonstrating the variety of ways that NbS is being conceptualised and operationalised in Australia.

The approaches mentioned most frequently (Table 2) were green infrastructure (n=24) urban greening (n=16), water sensitive urban design (n=14), urban green spaces (n=14), urban forests (n=11) and blue carbon sinks (n=9).

### 3.4 Problems addressed by NbS

A total of 54 problems were identified and grouped into nine overarching categories (Figure 5). Problems related to planning (n=71), water (n=61) and conservation (n=51) were the categories discussed most frequently (Figure 4). However, Adaptation, mitigation & resilience (n=22), urban planning (n=21) and biodiversity (n=17) were the most frequently mentioned key words in the literature on NbS in Australia (Figure 5).

**Fig. 5.**
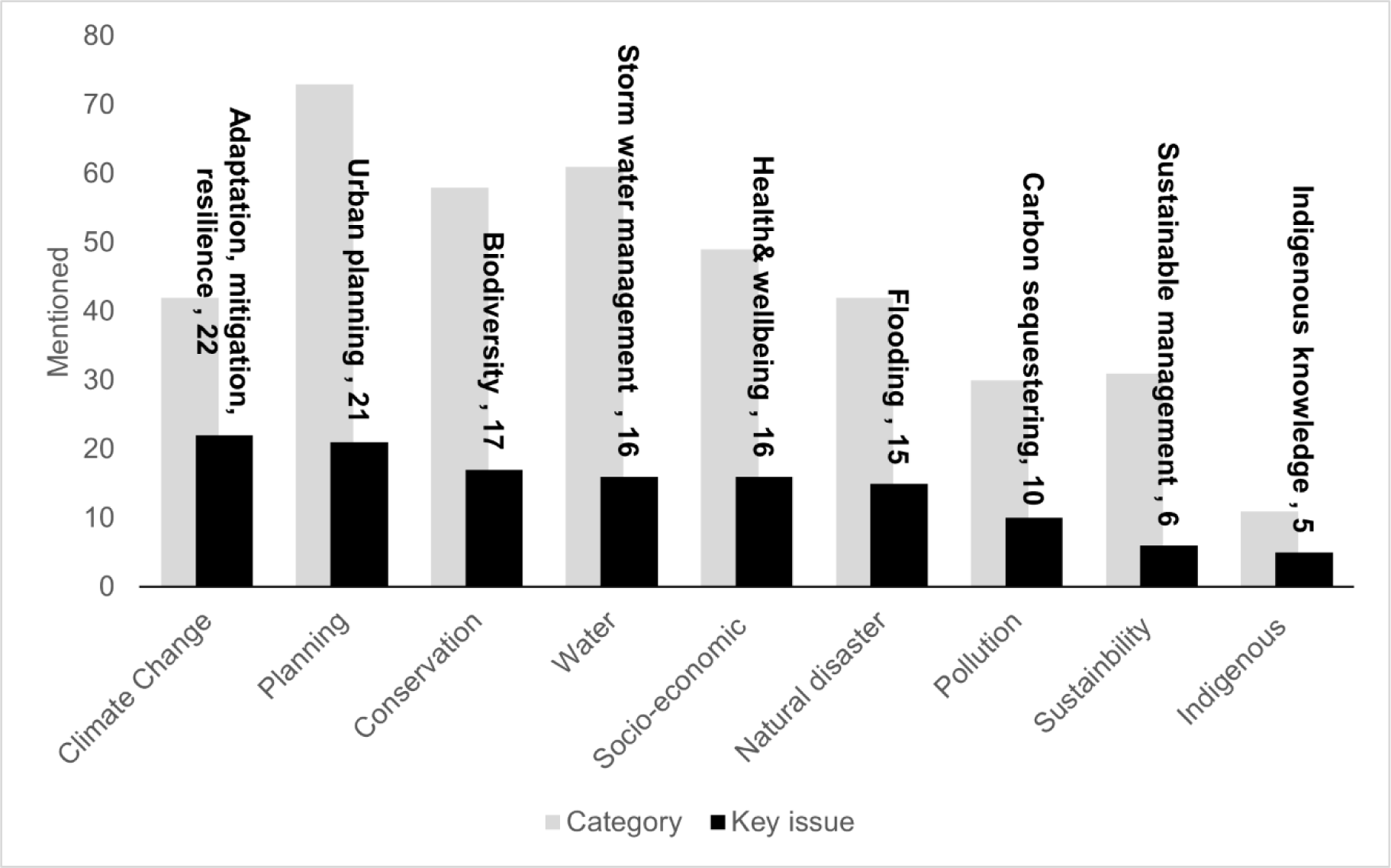
Key issues targeted by NbS in Australia. Total count of key words mentioned in each category sum greater than the number of papers reviewed as some papers discussed more than one issue in relation to NbS

Problems/issues discussed were primarily associated with land-based ecosystems. Urban (n=34) and terrestrial (n=14) ecosystems were discussed frequently, although agricultural ecosystems were only mentioned in Kingsley et al. (2021) and Ignatieva et al. (2020) (n=2). Comparatively, problems associated with NbS in coastal and freshwater ecosystems in Australia were mentioned far less than terrestrial topics (marine = 5, coastal = 8, rivers = 2, wetlands= 3). Some papers mentioned multiple ecosystems, meaning the total count of mentions may sum to more than the amount of papers reviewed.

### 3.5 Implementation of policies, strategies and frameworks

Of the policies, strategies and frameworks identified in the literature, three international frameworks were identified and mentioned the most frequently (n=21). The Paris Climate Agreement made up 42% of the total international mentions (n=16). Three federal and two state policies were also named (Table 3). Five local policies were identified and mentioned (n=11). For local strategies discussed, Melbourne was well-represented with the Living Melbourne Strategy (2019) being the most frequently mentioned (n=4) out of all Australian policies, and three different Melbourne strategies being mentioned cumulatively eight times in the literature (Table 3).

**Table 3.**
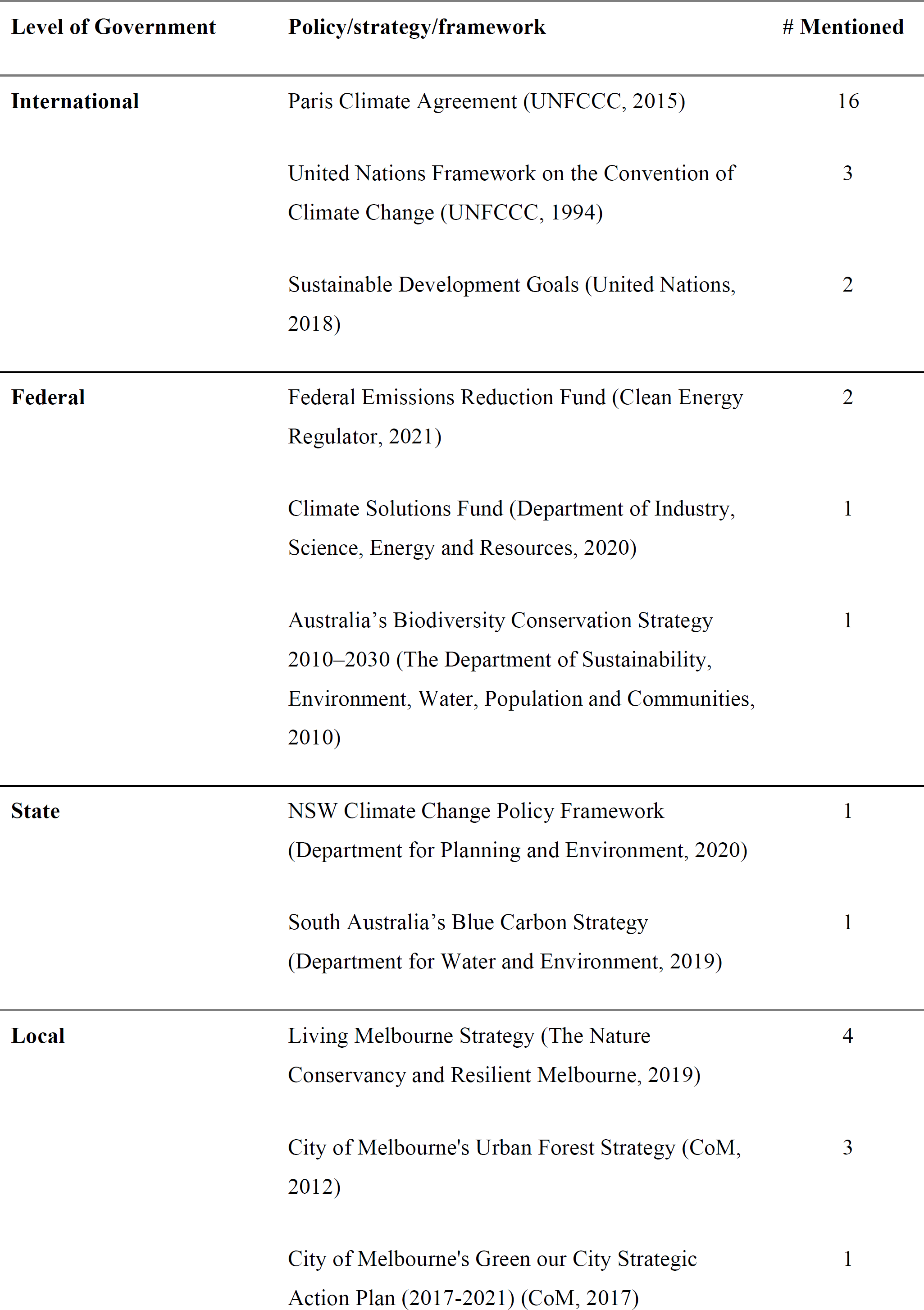
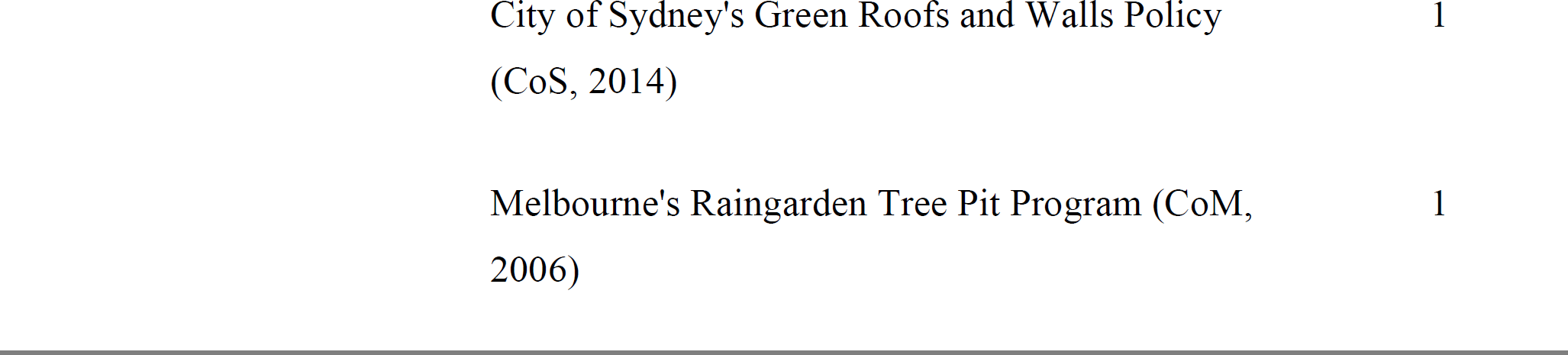
Policies, strategies, and frameworks identified in the literature. These are associated with applying the concept of NbS, but do not necessarily discuss NbS specifically.

| Level of Government | Policy/strategy/framework | # Mentioned |
| --- | --- | --- |
| **International** | Paris Climate Agreement (UNFCCC, 2015) | 16 |
|  | United Nations Framework on the Convention of Climate Change (UNFCCC, 1994) | 3 |
|  | Sustainable Development Goals (United Nations, 2018) | 2 |
| **Federal** | Federal Emissions Reduction Fund (Clean Energy Regulator, 2021) | 2 |
|  | Climate Solutions Fund (Department of Industry, Science, Energy and Resources, 2020) | 1 |
|  | Australia's Biodiversity Conservation Strategy 2010–2030 (The Department of Sustainability, Environment, Water, Population and Communities, 2010) | 1 |
| **State** | NSW Climate Change Policy Framework (Department for Planning and Environment, 2020) | 1 |
|  | South Australia's Blue Carbon Strategy (Department for Water and Environment, 2019) | 1 |
| **Local** | Living Melbourne Strategy (The Nature Conservancy and Resilient Melbourne, 2019) | 4 |
|  | City of Melbourne's Urban Forest Strategy (CoM, 2012) | 3 |
|  | City of Melbourne's Green our City Strategic Action Plan (2017-2021) (CoM, 2017) | 1 |

|  |  |
| --- | --- |
| City of Sydney's Green Roofs and Walls Policy<br>(CoS, 2014) | 1 |
| Melbourne's Raingarden Tree Pit Program (CoM,<br>2006) | 1 |

### 3.6 Scale of Implementation

Many articles identified local implementation (n=21), while regional/state-wide implementation (n=4) and national implementation (n=8) were less frequently discussed. Some articles did not discuss implementation at all (n=10).

### 3.7 Barriers

Articles often mentioned more than one barrier to implementation or recommendation to improve implementation. A total of 42% of articles recognised barriers when implementing NbS into policy (18). The most frequently mentioned barriers included lack of research and data (n=11), lack of resources (n=6), conflicting interests (n=6), lack of policy and frameworks (n=5) and lack of collaborative governance (n=5). A total of 28% of all articles gave recommendations on how to improve systems for implementing NbS (n=12). The most frequently mentioned recommendations included more research (n=16), policy improvements (n=13), increased engagement between levels of government (n=12), and increased collaboration across industry, research and government sectors (n=7). Despite eight articles discussing national implementation, none discussed the barriers to implementation at the federal scale. Barriers to implementation at a local (n=9) and state (n=5) scale were discussed more frequently.

## 4. Discussion

We found 43 studies exploring NbS in an Australian context (Appendix 1). These covered various topics and disciplines, including climate change, urban planning, conservation, sustainability, socio-economic, indigenous peoples, pollution, and natural disaster risk management. Perhaps surprisingly, definitions for NbS were scarce and varied between articles. Approaches (based on IUCN categories) to NbS were also varied and were being applied mostly in major cities, in particular Melbourne and Sydney (Maller, 2021; Kuller et al. 2021; Frantzeskaki & Bush 2021). The most common problems addressed by the NbS fell under the categories of climate change, urban planning, and biodiversity conservation (Frantzeskaki & Bush 2021; Coenen et al. 2020; Kirk et al. 2021). The scale of implementation was mostly local. Barriers to incorporating NbS into policy in Australia included a lack of research and data, lack of resources, conflicting interests, lack of policy and frameworks and lack of collaborative governance (Moosavi, Browne & Bush 2021).

The majority of the authors discussed the concept of NbS without providing a clear definition. There was no singular, comprehensive definition used for NbS in the literature. There is also a lack of guidance at the national level; Australia’s Strategy for Nature 2019-2030 (Commonwealth of Australia 2019) acknowledges how NbS is ‘critical to build the resilience of our unique nature’ but does not provide a definition of NbS. This lack of clarity of what NbS are and how they can help solve social-ecological problems in the Australian context may be hindering understanding, implementation and uptake of NbS in Australia. Although many papers do not define NbS specifically, the terms used to describe approaches to NbS provided context for what NbS means in Australia. Terms that are infrastructure-related, such as *urban greening* (Berthon, Thomas & Bekessy 2021), *water sensitive urban design* (Moosavi, Browne & Bush 2021), *urban green spaces* (Escobedo et al. 2019), *urban forests* (Esperon-Rodriguez et al. 2022), and an ecosystem-related protection/restoration term *blue carbon ecosystems* (Friess et al. 2020), are being used to refer to NbS in Australia the most frequently (Table 2). However globally accepted NbS terminology, as defined first by the IUCN (2020) and more recently revised and adopted by the United Nations Environment Assembly, has yet to be fully adopted in the Australian literature (Bush, Coffey & Fastenrath 2020; Uebel et al. 2021; Castonguay et al. 2018; Coenen et al. 2020; Pineda-Pinto et al. 2021; Wang et al. 2022). As a result, projects, research and studies which may technically be considered NbS may not use that terminology (Zhang et al. 2019; Greenway 2017 and Benites & Osmond 2021).

Internationally, competing definitions have been proposed by the IUCN, WWF and Oxford. Recently, the United Nations Environment Assembly (UNEA) has multilaterally agreed to a definition of nature-based solutions, based on the IUCN definition, that should focus future policy and research. This includes a definition of nature (‘natural or modified terrestrial, freshwater, coastal and marine ecosystems’), actions (‘to protect, conserve, restore, sustainably use and manage’ nature), problems to be addressed (‘social, economic and environmental challenges’) and the outcomes sought (‘simultaneously providing human well-being, ecosystem services and resilience and biodiversity benefits’). Australia has both a long tradition of ecosystem conservation in endemic, remnant ecosystems seeking improved biodiversity outcomes, and a large urban population who have benefits from urban green spaces and green infrastructure. Using the UNEA definition, NbS offer the possibility of bridging these approaches to ensure that both human wellbeing and biodiversity are considered in all nature-based policy and projects.

Adaptation, mitigation and resilience to climate change were the most frequently discussed challenges addressed by NbS. This was being led by international and local policy, with 37% of the papers discussing the Paris Climate Agreement. Local governments are using urban NbS in policy to help minimize adverse effects of climate change in cities and urban areas (Bayulken et al. 2021; Bush et al. 2020; Coenen et al. 2020; Croeser et al. 2020; Fastenrath et al. 2020; Frantzeskaki and Bush 2021; Gulsrud et al. 2018; Ordonez 2019; Pineda-Pinto et al. 2021; Zhang et al. 2019). For that reason, NbS for adaptation and effective climate policy is occuring from the “bottom-up” in Australia (Rayner 2010; Figure 6). In comparison, mention of state or federal policy, such as the Federal Emissions Reduction Fund, was scarce, highlighting Australia’s relatively slow uptake of NbS and historic inaction on climate change at a national level. Recent policies and a change in federal government may change this dynamic, e.g. Australia’s Strategy for Nature 2019-2030 (n=0) and the National Climate Resilience and Adaptation Strategy (DAWE 2021) (n=0) are both relevant to NbS.

**Fig. 6.**
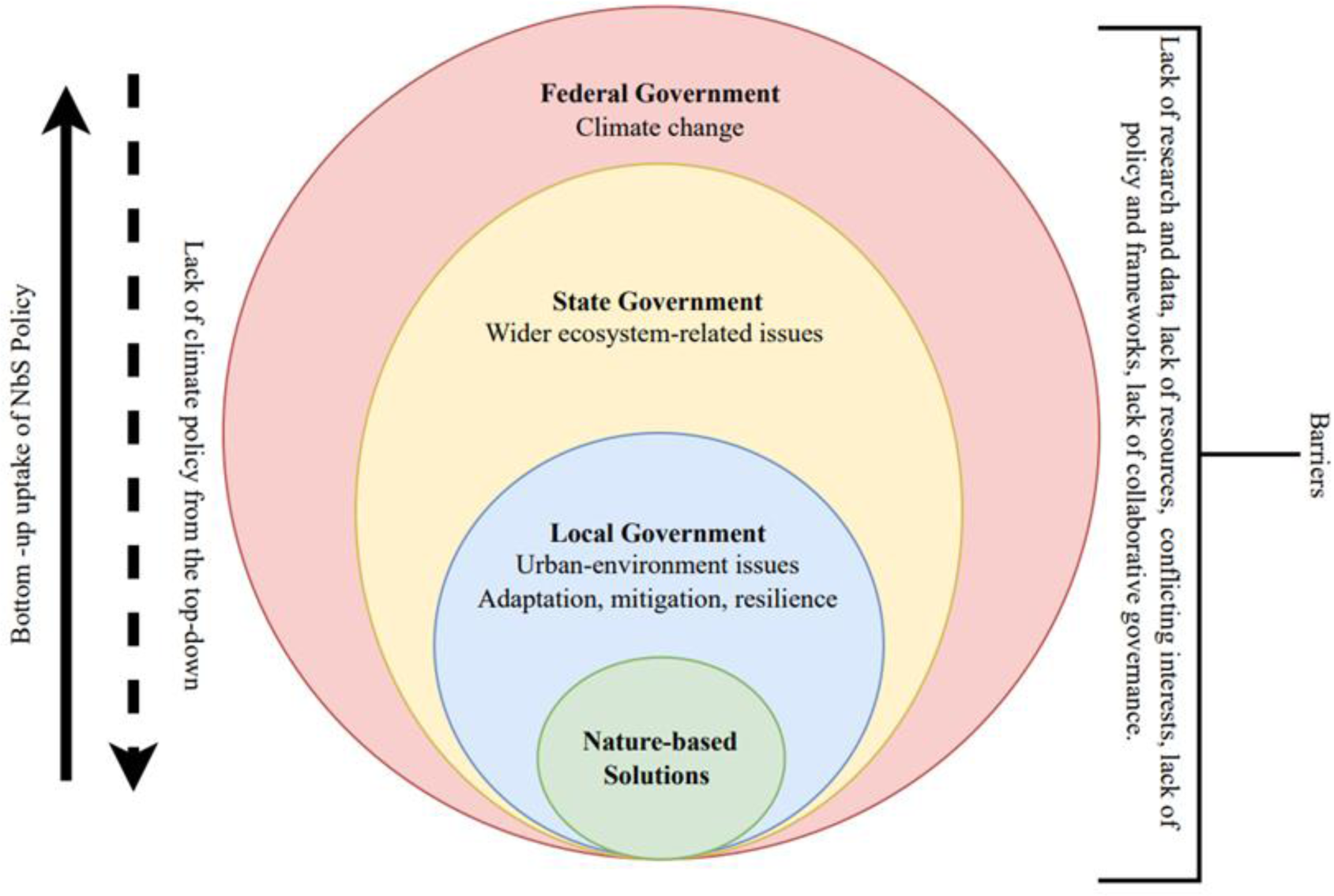
A conceptualisation of how NbS policy is used to address climate change issues in Australia and the barriers to its use

Bottom-up uptake of policy is emphasised by the number of local policies and frameworks that support NbS in Australia. The role of NbS for climate change adaptation “should be carefully designed and implemented through a bottom-up and participatory approach” in order to involve multiple stakeholders (Chausson et al. 2020). This is demonstrated in the urban planning strategies from Melbourne, such as the Living Melbourne Strategy (The Nature Conservancy and Resilient Melbourne, 2019), City of Melbourne’s Urban Forest Strategy (CoM 2012) and City of Melbourne’s Green Our City Strategic Action Plan 2017-2021 (CoM 2017), that were frequently discussed in the literature reviewed. Although this finding does not necessarily indicate the success of the strategies, it does show that these strategies are being referenced frequently in the discourse on NbS in Australia. The City of Melbourne’s Green Our City Strategic Action Plan 2017-2021 (CoM 2017) aims to address climate change issues that are compounded by urbanisation by implementing urban NbS. There is a particular focus on leading by example to expand urban greening in public areas, making relevant information and data available to the public, and introducing changes to the planning scheme. By demonstrating how urban planning can be used to solve ecosystem-wide challenges that exist in a human-environment interface, the City of Melbourne is leading the way in terms of implementing policy for NbS in Australia.

The conservation of biodiversity was another key issue mentioned frequently. Kirk et al. (2021) discuss the importance of biodiversity in sustaining human health and well-being in cities by looking at Biodiversity Sensitive Urban Design (BSUD) at Fisherman’s Bend in Melbourne, the largest urban renewal project in Australia. BSUD integrates targets for biodiversity into urban planning and development to “have a net benefit to native species and ecological communities through the provision of essential habitat and food resources’’ (Kirk et al. 2021; Benites and Osmond 2021; Berthon Thomas and Bekessy 2021). Similarly, ‘water sensitive urban design’ was discussed frequently as an approach to NbS occurring in Australia. This concept first emerged in Australia during the 1990’s and since then has been applied by practitioners around the world (Moosavi, Brown and Bush 2021). The City of Melbourne and The City of Sydney provide important lessons on how cities can transition towards water sensitive systems, particularly in stormwater management and coastal erosion (Kuller et al. 2021; Morris et al. 2019). Moosavi et al. (2021) points out that water management involves addressing drought and flood problems using NbS across multiple Australian states.

Although blue carbon ecosystems were identified frequently in the literature reviewed, actual applications and examples of water-based carbon-sequestration in Australia were discussed far less (Young et al. 2021). Blue carbon ecosystems, including vegetated coastal ecosystems such as tidal marshes, mangrove forests, and seagrass meadows, are protected and restored to sequester carbon from the atmosphere otherwise known as a “carbon sink” (Young et al. 2021). The only instance of blue carbon ecosystems in policy was the Blue Carbon Strategy (DAWE 2019) which aims to accelerate action to protect and restore coastal ecosystems in South Australia (Friess et al. 2020). Blue carbon ecosystems were often being proposed as policy solutions to meet climate change mitigation targets as they can be used to balance net emissions and reduce greenhouse gas emissions (Young et al. 2021; Morris et al. 2019; Shumway et al. 2021).

Perhaps unsurprising for the academic literature, lack of research and data was identified as a major barrier and more research identified as a key way to overcome this barrier. In comparison with other well established ecosystem management approaches such as biodiversity conservation (https://www.sciencedirect.com/science/article/pii/S2351989422002748) and urban greening (https://iopscience.iop.org/article/10.1088/1748-9326/abc5e4), there is indeed relatively little Australian research on NbS. As much of the thinking behind NbS occurred in Europe and North America, future research could develop a greater understanding of the particularities of the Australian context, particularly its unique flora and fauna, unevenly distributed human populations, large climatic gradients and tens of thousands of years of Indigenous culture and caring for country. NbS is already embedded in local government policy and practice, and bridging research with this existing policy and programs could be informative for both research and practice. Additional research and policy at sub-national and national scales could inform a key challenge for NbS – to scale up to effectively address large-scale challenges.

## 5. Conclusions and Recommendations

Ultimately, confusion around NbS definitions and terminology creates barriers to incorporating NbS into policy and practice. Definitions are an important tool in policy making and implementation as they provide clarity on what is included within the scope of policy and how policies can be implemented in practice (Cashore & Howlett 2020). International agreement now provides a clear definition of what problems NbS can address and what constitutes an effective NbS. The UNEA definition of NbS;

> *“actions to protect, conserve, restore, sustainably use and manage natural or modified terrestrial, freshwater, coastal and marine ecosystems, which address social, economic and environmental challenges effectively and adaptively, while simultaneously providing human well-being, ecosystem services and resilience and biodiversity benefits”* (UNEA 2022)

and the following eight principles from the IUCN would assist in defining NbS project and purpose in Australia (edited by Cohen-Shacham et al. 2016):

1. “*Embraces nature conservation norms (and principles);*
2. *can be implemented alone or in an integrated manner with other solutions to societal challenges (e.g. technological and engineering solutions);*
3. *are determined by site-specific natural and cultural contexts that include traditional, local and scientific knowledge;*
4. *produce societal benefits in a fair and equitable way, in a manner that promotes transparency and broad participation;*
5. *maintain biological and cultural diversity and the ability of ecosystems to evolve over time;*
6. *are applied at a landscape scale;*
7. *recognise and address the trade-offs between the production of a few immediate economic benefits for development, and future options for the production of the full range of ecosystems services; and*
8. *are an integral part of the overall design of policies, and measures or actions, to address a specific challenge.”* (IUCN 2020)

Improving engagement between the three levels of government, research institutions, industry and community will also be essential for NbS governance and policy development (Frantzeskaki & Bush 2021). The federal government could recognise and support the current infrastructure-related urban approaches to NbS projects occurring at a local level. Local urban planning strategies such as water-sensitive and biodiversity-sensitive urban design are addressing compounding socio-ecological challenges brought on by climate change and need to be included in federal budgets for NbS. Providing guidance on how to integrate NbS into climate adaption policy, like Chausson et al. (2020), would be beneficial in summarising the NbS options that can contribute to climate adaptation and sustainable development in Australia. Furthermore, additional research is needed on blue carbon approaches in Australia to ensure these are effectively benefiting human-well-being and biodiversity (thereby meeting the IUCN definition of NbS) and not merely acting as a “band-aid” policy for inaction on climate change.

## Acknowledgements

This project is supported with funding from the Australian Government’s National Environmental Science Program. We also acknowledge the Traditional Owners of Country throughout Australia and their continuing connection to land, sea, and community. We pay our respects to Traditional Owners, their cultures, and their Elders past, present and emerging.

## 7. Conflicts of interest

Author DK runs a consulting business, Future in Nature, which has received funding from the City of Melbourne related to urban forest policies (urban forest strategy) and programs (urban forest precinct planning) referred to in the manuscript. Authors DK and EJF have received funding from the Australian Government’s National Environmental Science Program through the Sustainable Communities and Waste Hub for nature-based solutions research; the primary research user for this Hub, with whom the research was codesigned, is the Department of Climate Change, Energy, the Environment and Water, which is responsible for Australia’s Strategy for Nature referred to in the article.

## Notes

### Competing Interest Statement

The authors have declared no competing interest.

