## Appendix 1 for "Nature-based solutions in Australia: a systematic quantitative literature review of terms, application and policy relevance"

Appendix 1. Nature-based Solution database

| ID | Title | Year | Authors | Journal |
| --- | --- | --- | --- | --- |
| 1 | How are nature-based solutions helping in the greening of cities in the context of crises such as climate change and pandemics? A comprehensive review | 2021 | Bayulken, Bogachan; Huisingh, Donald; Fisher, Peter M. J. | *Journal of Cleaner Production* |
| 2 | Bioconnections as Enablers of Regenerative Circularity for the Built Environment | 2021 | Benites, Henrique Sala; Osmond, Paul | *Urban Planning* |
| 3 | The role of ‘nativeness' in urban greening to support animal biodiversity | 2021 | Berthon, Katherine; Thomas, Freya; Bekessy, Sarah | *Landscape and Urban Planning* |
| 4 | Governing urban greening at a metropolitan scale: an analysis of the Living Melbourne strategy | 2020 | Bush, Judy; Coffey, Brian; Fastenrath, Sebastian | *Australian Planner* |
| 5 | Integrated modelling of stormwater treatment systems uptake | 2018 | Castonguay, A. C.; Iftekhar, M. S.; Urich, C.; Bach, P. M.; Deletic, A. | *Water Research* |
| 6 | Metropolitan governance in action? Learning from metropolitan Melbourne's urban forest strategy | 2020 | Coenen, Lars; Davidson, Kathryn; Frantzeskaki, Niki; Grenfell, Maree; Hakansson, Irene; Hartigan, Martin | *Australian Planner* |
| 7 | Patterns of tree removal and canopy change on public and private land in the City of Melbourne | 2020 | Croeser, Thami; Ordonez, Camilo; Threlfall, Caragh; Kendal, Dave; van der Ree, Rodney; Callow, David; Livesley, Stephen J. | *Sustainable Cities and Society* |
| 8 | Urban forests, ecosystem services, green infrastructure and nature-based solutions: Nexus or evolving metaphors? | 2019 | Escobedo, Francisco J.; Giannico, Vincenzo; Jim, C. Y.; Sanesi, Giovanni; Lafortezza, Raffaele | *Urban Forestry & Urban Greening* |
| 9 | Assessing climate risk to support urban forests in a changing climate | 2022 | Esperon-Rodriguez, Manuel; Rymer, Paul D.; Power, Sally A.; Barton, David N.; Carinanos, Paloma; Dobbs, Cynnamon; Eleuterio, Ana Alice; Escobedo, Francisco J.; Hauer, Richard; Hermy, Martin; Jahani, Ali; Onyekwelu, Jonathan C.; Ostberg, Johan; Pataki, Diane; R; rup, Thomas B.; Rasmussen, Torres; Roman, Lara A.; Russo, Alessio; Shackleton, Charlie; Solfjeld, Ingjerd; Doorn, Natalie S.; Wells, Matthew J.; Wistrom, | *Plants, People, Planet* |
| 10 | Scaling-up nature-based solutions. Lessons from the Living Melbourne | 2020 | Fastenrath, Sebastian; Bush, Judy; Coenen, Lars | *Geoforum* |

Appendix Table 4. Nature-based Solution database (*Continued)*

| ID | Title | Year | Authors | Journal |
| --- | --- | --- | --- | --- |
|  | strategy Melbourne, Australia |  |  |  |
| 11 | Governance of nature-based solutions through intermediaries for urban transitions – A case study from | 2021 | Frantzeskaki, Niki; Bush, Judy | *Urban Forestry & Urban Greening* |
| 12 | Mangrove blue carbon in the face of deforestation, climate change and restoration | 2020 | Friess, Daniel A.; Krauss, Ken W.; Taillardat, Pierre; Adame, Maria Fern; a; Y; o, Erik S.; Cameron, Clint; Sasmito, Sigit D.; Sillanpaa, Meriadec | *Annual Plant Reviews* |
| 13 | Stormwater wetlands for the enhancement of environmental ecosystem services: case studies for two retrofit wetlands in Brisbane, Australia - ScienceDirect | 2017 | Greenway, Margerate | *Journal of Cleaner Production* |
| 14 | Innovative urban forestry governance in Melbourne?: Investigating “green placemaking” as a nature-based solution | 2018 | Gulsrud, Natalie Marie; Hertzog, Kelly; Shears, Ian | *Environmental Research* |
| 15 | What makes a successful Sponge City project? Expert perceptions of critical factors in integrated urban water management in the Asia-Pacific | 2021 | Hawken, Scott; Sepasgozar, S. M. E.; Prodanovic, Veljko; Jing, Jia; Bakelmun, Ashley; Che, Shengquan; Zhang, Kefeng | *Sustainable Cities and Society* |
| 16 | Contemporary Oyster Reef Restoration: Responding to a Changing World | 2021 | Howie, Alice H.; Bishop, Melanie J. | *Frontiers in Ecology and Evolution* |
| 17 | Lawns in Cities: From a Globalised Urban Green Space Phenomenon to Sustainable Nature-Based Solutions | 2020 | Ignatieva, Maria; Haase, Dagmar; Dushkova, Diana; Haase, Annegret | *Land* |
| 18 | Reshaping forest management in Australia to provide nature-based solutions to global challenges | 2021 | Jackson, W.' Freeman, M; Freeman, B.; Parry-Husbands, H | *Australian Forestry* |
| 19 | Urban agriculture as a nature-based solution to address socio-ecological challenges in Australian Cities | 2021 | Kingsley, Jonathan; Egerer, Monika; Nuttman, Sonia; Keniger, Lucy; Pettitt, Philip; Frantzeskaki, Niki; Gray, Tonia; Ossola, Alessandro; Lin, Brenda; Bailey, Aisling; Tracey, Danielle; Barron, Sara; Marsh, Pauline. | *Urban Forestry &*  *Urban Greening* |
| 20 | Building biodiversity into the urban fabric: A case study in applying Biodiversity Sensitive Urban Design (BSUD) | 2021 | Kirk, Holly; Garrard, Georgia E.; Croeser, Thami; Backstrom, Anna; Berthon, Katherine; Furlong, Casey; Hurley, Joe; Thomas, Freya; Webb, Anissa; Bekessy, Sarah A. | *Urban Forestry & Urban Greening* |

Appendix Table 4. Nature-based Solution database *(Continued)*

| ID | Title | Year | Authors | Journal |
| --- | --- | --- | --- | --- |
| 21 | A planning-support tool for spatial suitability assessment of green urban stormwater infrastructure | 2019 | Kuller, Martijn; Bach, Peter M.; Roberts, Simon; Browne, Dale; Deletic, Ana | *Science of Total Environment* |
| 22 | Are we planning blue-green infrastructure opportunistically or strategically? Insights from Sydney, Australia | 2021 | Kuller, Martijn; Reid, David J; Prodanovic, Veljko | *Blue-Green Systems* |
| 23 | Designing collaborative governance for nature-based solutions | 2021 | Malekpour, Shirin; Tawfik, Sylvia; Chesterfield, Chris | *Urban Forestry & Urban Greening* |
| 24 | Re-orienting nature-based solutions with more-than-human thinking | 2021 | Maller, Cecily | *Cities* |
| 25 | Perceptions of nature-based solutions for Urban Water challenges: Insights from Australian researchers and practitioners | 2021 | Moosavi, Sareh; Browne, Geoffrey R.; Bush, Judy | *Urban Forestry & Urban Greening* |
| 26 | Developing a nature-based coastal defence strategy for Australia | 2019 | Morris, RebeccaL.; Strain, Elisabeth M. A.; Konlechner, Teresa M.; Fest, Benedikt J.; Kennedy, David M.; Arndt, Stefan K.; Swearer, Stephen E. | *Australian Journal of Civil Engineering* |
| 27 | Understanding socio-economic benefits of stormwater management system through urban lakes in Western Sydney, Australia | 2018 | Natarajan, Sai Kiran; Hagare, Dharmappa; Maheshwari, Basant | *Ecohydrology & Hydrobiology* |
| 28 | International approaches to protecting and retaining trees on private urban land | 2021 | Ordonez-Barona, Camilo; Bush, Judy; Hurley, Joe; Amati, Marco; Juhola, Sirkku; Frank, Stephen; Ritchie, Myles; Clark, Christopher; English, Alex; Hertzog, Kelly; Caffin, Meg; Watt, Steve; Livesley, Stephen J. | *Journal of Environmental Management* |
| 29 | Polycentric governance in nature-based solutions: insights from Melbourne urban forest managers | 2019 | Ordonez, Camilo | *Landscape Architecture Frontiers* |
| 30 | How Urban Forest Managers Evaluate Management and Governance Challenges in Their Decision-Making | 2020 | Ordonez, Camilo; Kendal, Dave; Threlfall, Caragh G.; Hochuli, Dieter F.; Davern, Melanie; Fuller, Richard A.; van der Ree, Rodney; Livesley, Stephen J. | *Forests* |
| 31 | Making nature-based solutions  climate-ready for the 50 °C world | 2021 | Ossola, Alessandro; Lin, Brenda B. | *Environmental Science & Policy* |
| 32 | A case study balancing predetermined targets and real- | 2020 | Parker, Jackie; Simpson, Greg D. | *Forests* |

Appendix Table 4. Nature-based Solution database *(Continued)*

| ID | Title | Year | Authors | Journal |
| --- | --- | --- | --- | --- |
|  | world constraints to guide optimum urban tree canopy cover for Perth, Western Australia |  |  |  |
| 33 | Mapping social-ecological injustice in Melbourne, Australia: An innovative systematic methodology for planning just cities | 2021 | Pineda-Pinto, Melissa; Nygaard, Christian A.; Ch; rabose, Manoj; Frantzeskaki, Niki | *Land Use Policy* |
| 34 | Water Sensitive Cities Index: A diagnostic tool to assess water sensitivity and guide management actions | 2020 | Rogers, B. C.; Dunn, G.; Hammer, K.; | *Water Research* |
| 35 | Nature-Based Urbanization: Scan Opportunities, Determine Directions and Create Inspiring Ecologies | 2021 | Roggema, Rob; Tillie, Nico; Keeffe, Greg | *Land* |
| 36 | Policy solutions to facilitate restoration in coastal marine environments | 2021 | Shumway, Nicole; Bell-James, Justine; Fitzsimons, James A.; Foster, Rose; Gillies, Chris; Lovelock, Catherine E. | *Marine Policy* |
| 37 | Urban green space soundscapes and their perceived restorativeness | 2021 | Uebel, Konrad; Marselle, Melissa; Dean, Angela J.; Rhodes, Jonathan R.; Bonn, Aletta | *People and Nature* |
| 38 | Future climate impacts on forest growth and implications for carbon sequestration through reforestation in southeast Australia | 2022 | Wang, Bin; Waters, Cathy; Anwar, Muhuddin Rajin; Cowie, Annette; Li Liu, De; Summers, David; Paul, Keryn; Feng, Puyu | *Journal of Environmental Management* |
| 39 | Ten years of greening a wide brown land: A synthesis of Australian green roof research and roadmap forward | 2021 | Williams, Nicholas S. G.; Bathgate, Rachael S.; Farrell, Claire; Lee, Kate E.; Szota, Chris; Bush, Judy; Johnson, Katherine A.; Miller, Rebecca E.; Pianella, Andrea; Sargent, Leisa D.; Schiller, Julia; Williams, Kathryn J. H.; Rayner, John P. | *Urban Forestry & Urban Greening* |
| 40 | Urban green roofs promote metropolitan biodiversity: A comparative case study | 2022 | Wooster, E. I. F.; Fleck, R.; Torpy, F.; Ramp, D.; Irga, P. J. | *Building and Environment* |
| 41 | National scale predictions of contemporary and future blue carbon storage | 2021 | Young, Mary A.; Serrano, Oscar; Macreadie, Peter, I; Lovelock, Catherine E.; Carnell, Paul; Ierodiaconou, Daniel | *Science of the Total Environment* |
| 42 | Evaluating the reliability of stormwater treatment systems under various future climate conditions | 2019 | Zhang, Kefeng; Manuelpillai, Desmond; Raut, Bhupendra; Deletic, Ana; Bach, Peter M. | *Journal of Hydrology* |

Appendix Table 4. Nature-based Solution database *(Continued)*

| ID | Title | Year | Authors | Journal |
| --- | --- | --- | --- | --- |
| 43 | The effect of intermittent drying and wetting stormwater cycles on the nutrient removal performances of two vegetated biofiltration designs | 2021 | Zinger, Yaron; Prodanovic, Veljko; Zhang, Kefeng; Fletcher, Tim D.; Deletic, Ana | *Chemosphere* |
