## Appendix 2 for "Nature-based solutions in Australia: a systematic quantitative literature review of terms, application and policy relevance"

**Appendix 2. NbS terms used in Australia**

| IUCN Recognized Terms | Other Synonyms Used in Australia |
| --- | --- |
| Ecological restoration | Blue carbon sinks/ecosystems |
| Ecological engineering | Biodiversity Sensitive Urban Design |
| Forest landscape restoration | Water Sensitive Urban Design |
| Ecosystem- based adaptation | Urban forests |
| Ecosystem-based mitigation | Urban Greening |
| Climate adaptation services | Urban agriculture |
| Ecosystem- based disaster risk reduction | Nature-based coastal defence |
| Green infrastructure | Urban green design / green urbanism |
| Integrated coastal zone management | Reforestation |
| Integrated water resource management | (Urban) green spaces |
| Area-based conservation |  |
